# Inhibition of the NLRP3 Inflammasome Prevents Hyperglycemia in a Humanized Rodent Model of Type 2 Diabetes

**DOI:** 10.64898/2026.06.19.733338

**Authors:** Daniel T. Meier, Elise Dalmas, Leila Rachid, Audrey Le Guernic, Zora Baumann, Nicolas Venteclef, Marc Y. Donath

## Abstract

The NACHT, LRR and PYD domains-containing protein 3 (NLRP3) inflammasome is a protein complex that senses metabolic disturbances and in response processes IL-1beta. Prolonged activation of the NLRP3 inflammasome by metabolic stress induces chronic low-grade inflammation and contributes to the development type 2 diabetes and its comorbidities. Here, we developed a mouse model of type 2 diabetes that features impaired proinsulin processing and the ability to form islet amyloid plaques, two determinants of human type 2 diabetes pathology. We show that at advanced ages, these mice develop amyloidosis and inflammation in their insulin-producing pancreatic islets. Beta-cell mass and function were impaired in these mice and severe hyperglycemia developed within 6 months. Oral application of OLT1177, a selective inhibitor of the NLRP3 inflammasome, prevented the development of hyperglycemia. Our data show that inhibition of the NLRP3 inflammasome in a humanized mouse model of severe type 2 diabetes prevents the development of amyloid-associated hyperglycemia.

## Introduction

The NACHT, LRR and PYD domains-containing protein 3 (NLRP3) inflammasome complex is highly expressed in innate immune cells. It processes and releases the cytokines IL-1beta and IL-18 in response to sensing cellular stress signals. Inflammasome activators include obesity- and diabetes-associated elevated glucose, cholesterol, uric acid, non-esterified fatty acids, extracellular ATP and islet amyloid polypeptide (IAPP) [1–6]. Assembly of a functional protein complex which includes NLRP3, apoptosis-associated speck-like protein containing a caspase recruitment domain (ASC, *Pycard*) and caspase 1 then allows to process pro-IL-1beta and pro-IL18 and to release the active form of these pro-inflammatory cytokines. Prolonged activation of the NLRP3 inflammasome drives chronic low-grade inflammation, which is associated with the development of type 2 diabetes and associated secondary comorbidities [7].

The beta-cell product IAPP is a peptide hormone co-secreted with insulin [8] and may aggregate into plaques which are found as amyloid deposits in most patients with type 2 diabetes [9]. Oligomerization of IAPP is cytotoxic and the degree of amyloid deposition in the pancreatic islets is associated with beta-cell apoptosis [10] and inversely correlated with beta-cell mass and function [10, 11]. Oligomers of human IAPP triggers IL-1beta secretion via activation of the NLRP3 inflammasome [6] while rodent IAPP is non-amyloidogenic and not cytotoxic [12, 13]. Therefore, several rodent research models transgenically expressing human IAPP were developed to study amyloidosis. Interestingly, glucose may also trigger IL-1beta secretion [14] and both glucose [15] and non-esterified fatty acids [16] promote islet amyloidosis in human islets via upregulation of IAPP expression.

Type 2 diabetes is associated with increased numbers of macrophages in islets [17] and islet inflammation contributes to the pathology of type 2 diabetes [18, 19]. Islet amyloidosis is a major determinant of this inflammatory state [20]. Further, in human islet cultures, IL-1beta increases amyloid deposition [21]. Downstream of the NLRP3 inflammasome, prolonged IL-1beta exposure induces beta-cell dysfunction and induces insulin resistance, contributing to the development of type 2 diabetes [2, 22–24]. Consequently, blocking IL-1 signaling results in improved beta-cell function and glucose tolerance in both rodent models of islet amyloidosis [25, 26] as well as patients with type 2 diabetes [27, 28].

Inhibition of the NLRP3 inflammasome upstream of IL-1 signaling provides another entry point to target inflammatory states. Consequently, several NLRP3 inhibitors were developed and entered clinical evaluation [29]. The targeted diseases were of chronic highly inflammatory nature, including cryopyrin-associated periodic syndrome (CAPS), joint inflammation, gout, neurodegenerative diseases such as Alzheimer disease and Parkinson disease, ulcerative colitis and chronic obstructive pulmonary disease. More recently, research was expanded to chronic low-grade inflammatory pathologies such as obesity and diabetes and its associated cardiometabolic co-morbidities [30]. As such, the NLRP3 inhibitor VTX3232 was shown to prevent a variety of cardiometabolic parameters including hyperglycemia, glucose tolerance and lipid metabolism in diet-induced obese mice [31]. These changes were associated with reduced food intake and body weight gain which in addition to the anti-inflammatory properties of this compound might explain the observed metabolic improvements. A similar effect on body weight regulation and potentially secondary effects on metabolic parameters in high-fat diet-fed mice was observed with two more NLRP3 inhibitors, NT-0249 and NT-0796 [32].

OLT1177 (dapansutrile, 3-methanesulfonyl-propionitrile) is a small orally available brain-penetrating molecule that selectively inhibits the NLRP3 inflammasome [33]. Previous experimental models of infection, acute arthritis, myocardial infarction and experimental autoimmune encephalomyelitis showed the efficacy of this drug in reducing inflammation [33–36]. In humans, OLT117 was found to be safe and well tolerated [33]. In a proof-of-concept, phase 2a trial, OLT1177 was shown to reduce joint pain in patients with gouty arthritis [37]. In another study with patients with heart failure and type 2 diabetes, OLT1177 reduced fasting blood glucose values in a dose-dependent manner with concomitant improvements in cardiac function and exercise capacity [38].

Current preclinical models of type 2 diabetes are largely limited to diet-induced obesity models or genetic models with defects in leptin signaling, such as ob/ob and db/db mice. While valuable, these models do not fully recapitulate the human disease, which is characterized by islet amyloid deposition, impaired proinsulin processing, and progressive beta-cell dysfunction. Moreover, because NLRP3 inhibition has been shown to reduce obesity and improve metabolic parameters, the interpretation of results in obesity-driven models may be confounded by weight loss–related effects. To specifically address the role of the NLRP3 inflammasome in islet failure, we developed a mouse model that incorporates three key features of human type 2 diabetes: impaired proinsulin processing, the capacity to develop islet amyloidosis, and ageing, while avoiding the confounding influence of obesity. Using this model, we tested whether treatment with OLT1177 could prevent the development of beta-cell dysfunction and hyperglycemia through inhibition of the NLRP3 inflammasome.

## Results

### Enabling islet amyloid deposition in combination with impaired proinsulin processing induces hyperglycemia

The pathology of human type 2 diabetes includes the production and deposition of amyloid fibrils in pancreatic islets [9, 10, 19] and impaired proinsulin processing [39–41]. To generate a mouse model that includes these two features of human type 2 diabetes, we assessed mice that express human islet amyloid polypeptide (IAPP) in their pancreatic beta cells [42]. On a FVB/N genetic background, males showed mildly increased blood glucose values, although this did not reach statistical significance (Fig. 1A) while body weight was similar to control mice (Fig. 1B). Glycemia and body weight in female mice expressing human IAPP did not differ from controls (Fig. S1A-B). To model impaired proinsulin processing, we generated mice with beta cell-specific inducible ablation of prohormone convertase 1/3 (PC1/3). We previously reported that these mice develop hyperglycemia and obesity on a C57BL/6N genetic background [43]. However, ablation of PC1/3 in these mice on an FVB/N genetic background did not induce hyperglycemia but still lead to mild overweight (Fig. 1 C-E). Females had unchanged glycemia and body weight development (Fig. S1 C-D). We then combined these two mouse models and generated mice with inducible impairment of proinsulin processing and the ability to form amyloid plaques. In uninduced mice, body weight was similar between genotypes but mice carrying the human IAPP transgene were mildly hyperglycemic compared to mice expressing endogenous islet amyloid polypeptide only (Fig. S1 E-F). PC1/3 knockout induction at 8 weeks of age did not change glycemia but mice capable of forming amyloid plaques became frankly hyperglycemic, reaching glucose levels comparable to those observed in severely decompensated human type 2 diabetes at 28 weeks of age (Fig. 1 F-G). Body weight did not differ between genotypes (Fig. 1 H). These data show that carrying the transgene that expresses human IAPP induces mild hyperglycemia and that in combination with impairment of proinsulin processing and ageing, mice develop severe hyperglycemia.

**Figure 1.**
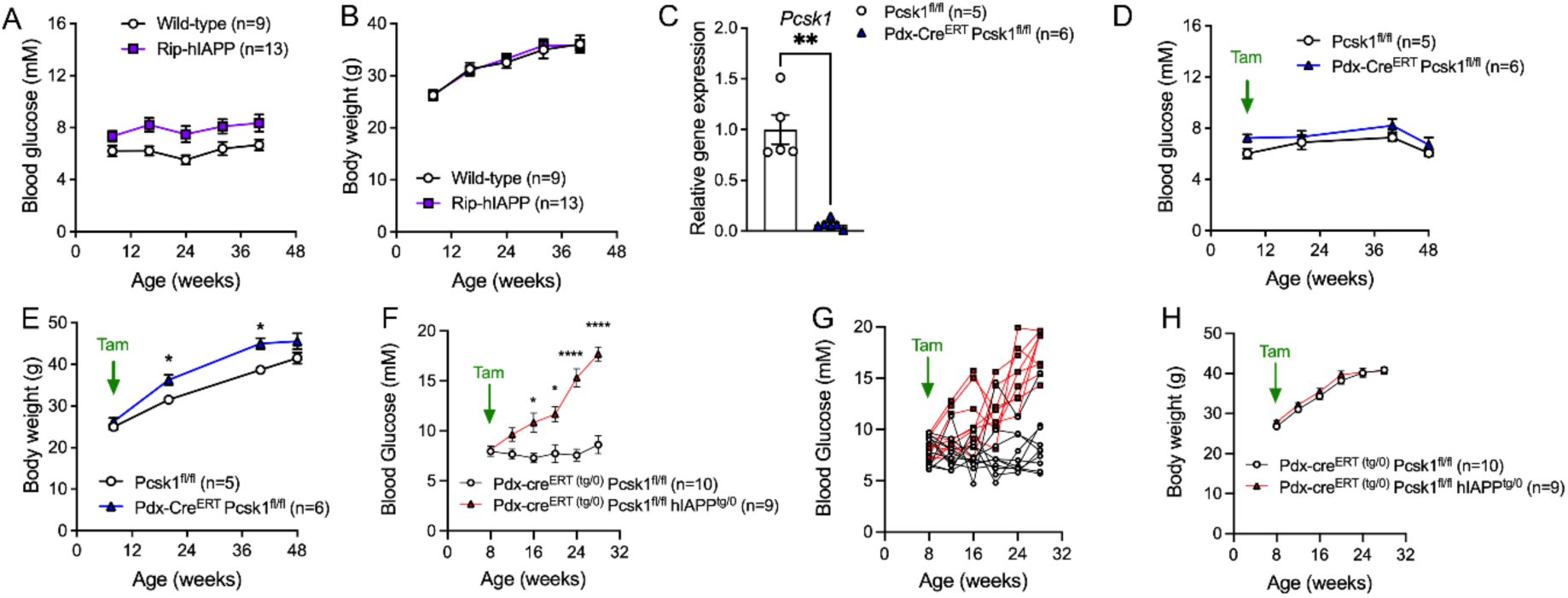
Enabling islet amyloid deposition in combination with impaired proinsulin processing induces hyperglycemia. (A) Glycemia and (B) body weight development in male hIAPP mice on an FVB/N genetic background. (C) Gene expression levels in islets isolated from 13-15-week-old Pdx-cre^ERT^ Pcsk1^fl/fl^ mice that had received tamoxifen at 8 weeks of age. (D) Glycemia and (E) body weight development in Pdx-cre^ERT^ Pcsk1^fl/fl^ mice that had received tamoxifen at 8 weeks of age. (F) Average glycemia per experimental group, (G) glycemia of individual mice and (H) body weight development of Pdx-cre^ERT^ Pcsk1^fl/fl^ hIAPP^tg/0^ mice that had received tamoxifen at 8 weeks of age. Statistics: A, B, D, E, F, H 2-way ANOVA with Sidak’s multi comparison test. C: Mann-Whitney test. * p<0.05, ** p< 0.01, ***p<0.001, ****p<0.0001.

### Amyloid deposition is associated with reduced beta-cell mass

Human type 2 diabetes is associated with the accumulation of cytotoxic amyloid plaques in the islets [9, 10], a feature that only mice carrying the human IAPP transgene have [42]. We therefore analyzed pancreata of Pdx-cre^ERT^ Pcsk1^fl/fl^ hIAPP mice histologically. As expected, mice with reduced PC1/3 expression but that only express endogenous mouse IAPP remained normoglycemic and did not show amyloid depositions. Littermates expressing human IAPP became hyperglycemic and all showed amyloid depositions (Fig. 2 A, Fig. S2 A-B). 88% of their islets showed amyloid deposits and on average 10% of their islet area covered with amyloid plaques and this was associated with a marked reduction in insulin-positive islet area (Fig. 2 B-D). Beta-cell mass was reduced which was based on reduced pancreas weight and smaller islets, while the number of islets was unchanged between genotypes (Fig. 2 E-H). These data show that Pdx-cre^ERT^ Pcsk1^fl/fl^ hIAPP mice develop islet amyloidosis which is associated with reduced beta-cell mass.

**Figure 2.**
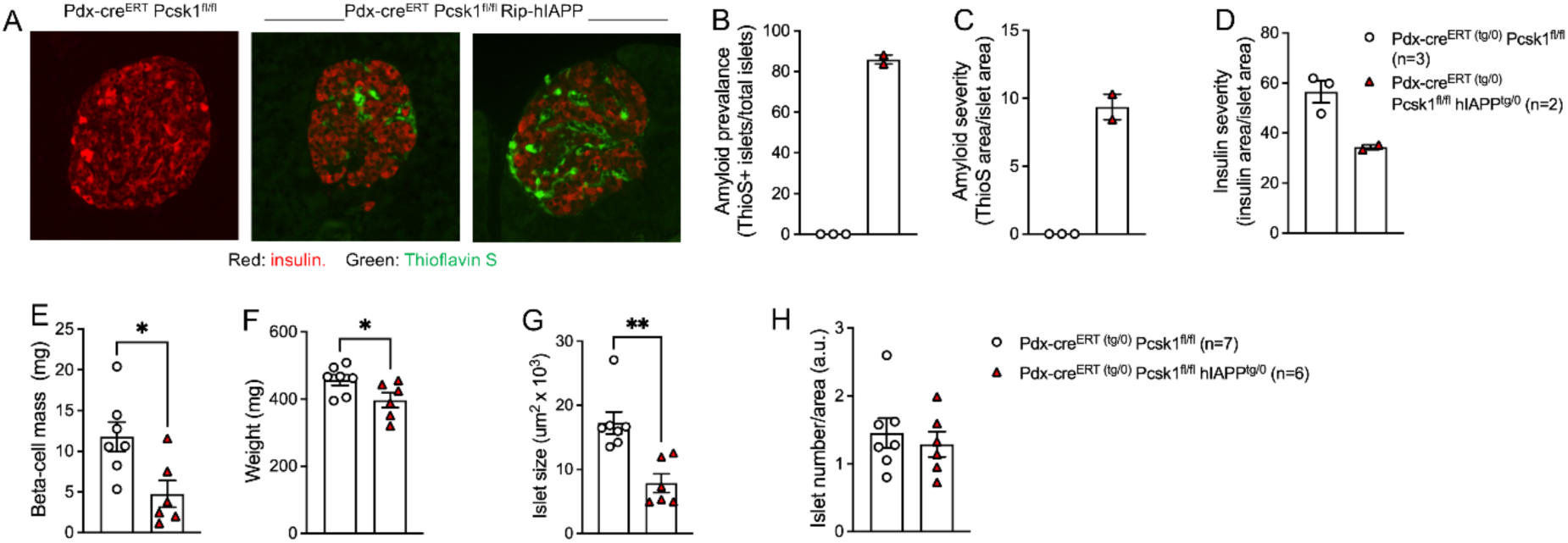
Amyloid deposition is associated with reduced beta-cell mass. (A) Immunohistochemical images of single islets in pancreatic sections from Pdx-cre^ERT^ Pcsk1^fl/fl^ hIAPP^tg/0^ mice stained for insulin (red) and amyloid (green). (B) Amyloid prevalence (% Thioflavin-S-positive islets), (C) amyloid severity (% Thioflavin-S-positive islet area) and (D) insulin severity (% insulin-positive area) in 29-week-old Pdx-cre^ERT^ Pcsk1^fl/fl^ hIAPP^tg/0^ mice. (E) Beta-cell mass, (F) pancreas weight, (G) average islet size and (H) islet number per area in Pdx-cre^ERT^ Pcsk1^fl/fl^ hIAPP^tg/0^ mice at sacrifice. Statistics: B-H: Mann-Whitney test. * p<0.05, ** p< 0.01.

### Amyloid deposition is associated with increased numbers of islet macrophages and islets inflammation

Next, we attempted to isolate pancreatic islets by collagenase digestion from hyperglycemic 32-week-old mice and repeated the procedure in 28-week-old mice, but were unable to obtain suitable islets from Pdx-cre^ERT^ Pcsk1^fl/fl^ hIAPP mice (Fig. S2C), potentially because the remaining islets were too fibrotic to be liberated from the tissue. We then isolated islets from 2 cohorts of 16-week-old mice that were mildly hyperglycemic (Fig. 3 A-B). The expression level of the chemokine *Cxcl1* (encoding KC) but not *Ccl2* (encoding MCP-1) was increased in mice expressing human IAPP (Fig. 3 C). The pan immune cell marker CD45 as well as macrophage marker Itgax (encoding Cd11c) were increased but the T cell marker *Cd3e* remained unchanged (Fig. 3 D). Expression of cytokines *Il1b* and *Il6* was increased, *Tnf* was unchanged and *Foxp3* and *Ms4a1*, markers for regulatory T cells and B cells, were below the detection limit (Fig. 3 E). Caspase-1 was increased while the expression level of other members of the IL-1 pathway (*Il1r1*, *Il1rn*, *Nlrp3* and *Pycard*) were not influenced by amyloidosis (Fig. 3 F). To confirm these findings on a protein level, we next quantified macrophage counts in pancreatic sections from a cohort of mice at sacrifice (Fig. S2 D). As observed in previous cohorts of mice, islet area and number was reduced (Fig. S2 E and Fig 3 G). The density of intra-islet as well as peri-islet macrophages was increased (Fig. 3 H-I), confirming RNA expression data. Next, we assessed whether this local inflammatory state in the pancreas can also be detected in the circulation. However, at 17 weeks of age, a timepoint when the specific cohort of mice showed considerable hyperglycemia, circulating concentrations of myeloid cells, neutrophils and T cells did not differ between genotypes (Fig. S2 F-I). These data show that islet amyloidosis is associated with increased number of pancreatic islet macrophages and expression of *Il1b* and *Il6*.

**Figure 3.**
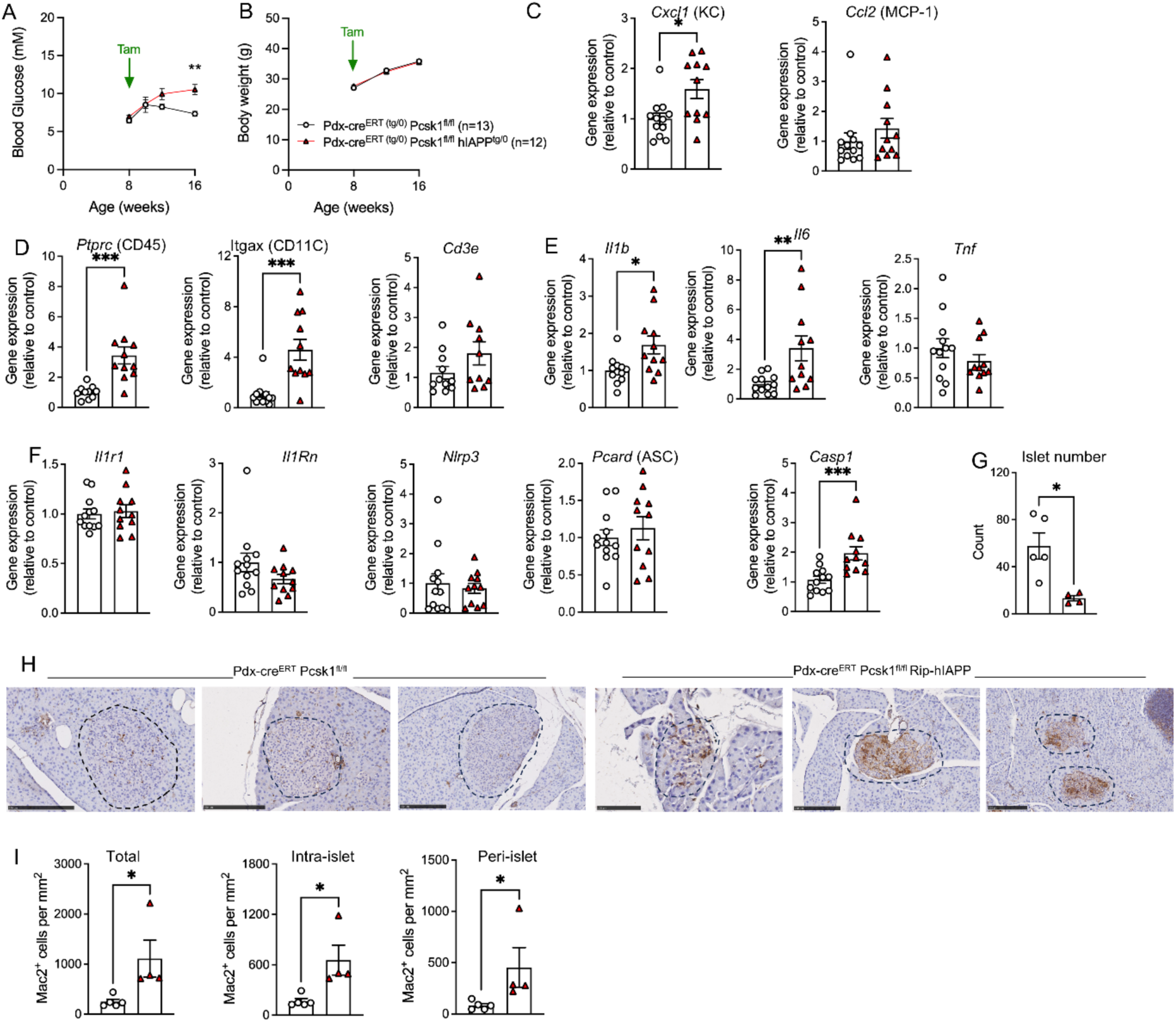
Amyloid deposition is associated with increased numbers of islet macrophages and islets inflammation. (A) Glycemia and (B) body weight development of a cohort of Pdx-cre^ERT^ Pcsk1^fl/fl^ hIAPP^tg/0^ mice used for islet isolation. (C) Chemokine, (D) immune cell, (E) cytokine and (F) IL-1-related transcripts in islets isolated from 16-week-old Pdx-cre^ERT^ Pcsk1^fl/fl^ hIAPP^tg/0^ mice. (G) Islet number per section, (H) representative Mac2 staining of pancreas sections and (I) quantification of Mac2 staining in pancreas section from 28-week-old Pdx-cre^ERT^ Pcsk1^fl/fl^ hIAPP^tg/0^ mice. Statistics: A 2-way ANOVA with Sidak’s multi comparison test. C-G, I: Mann-Whitney test. * p<0.05, ** p< 0.01, ***p<0.001.

### The NLRP3 inhibitor OLT1177 prevents the development of islet amyloid-induced hyperglycemia

Since islet amyloid activates the NLRP3 inflammasome to induce IL-1β secretion and islet inflammation [6], we hypothesized that inhibition of this pathway might prevent beta-cell failure in our mouse model of type 2 diabetes. To test this hypothesis, PC1/3 knockout in Pdx-cre^ERT^ Pcsk1^fl/fl^ hIAPP mice was induced at 8 weeks of age and mice were fed a diet containing OLT1177 (dapansutrile; 3-methanesulfonyl-propionitrile), a small orally available selective NLRP3 inflammasome inhibitor [33], starting at 9 weeks of age. Mice receiving control diet developed severe hyperglycemia while animals fed OLT1177 had blood glucose values comparable to mice unable to form amyloid deposits (Fig. 4 A). One cohort of mice had to be sacrificed at 20 weeks of age due metabolic decompensation of the control group and the remaining mice were sacrificed at 26 weeks of age (Fig. 4 B). Mice fed control diet but not OLT1177-fed mice had increased peak glycemia during the study compared to mice unable to form amyloid deposits (Fig. 4 C). These results align well with a human intervention study in which the subgroup of patients with type 2 diabetes showed a 4.6-fold decrease in fasting blood glucose compared to placebo after 14 days of OLT1177 treatment [38] (Supplemental Table 5). Body weight was comparable between genotypes and not influenced by OLT1177 (Fig. 4 D). One potential mechanism how inhibition of the NLRP3 inflammasome might improve glycemia is via reducing IL-1beta-induced insulin secretion [44], a concept termed “beta-cell rest” [30]. To test whether reducing the workload of insulin-producing cells improves glycemia in our mouse model of type 2 diabetes, we next fed Pdx-cre^ERT^ Pcsk1^fl/fl^ hIAPP mice a diet containing the potassium channel opener diazoxide which limits insulin secretion. Amyloid mice fed a control diet became overtly hyperglycemic and some had to be taken out of the experiment due to severe metabolic decompensation (Fig. 4 E-F). Amyloid mice receiving diazoxide also showed mildly increased glycemia compared to non-amyloid mice, but the highest value measured during the study was 39% lower in mice receiving diazoxide (Fig. 4 G). Body weight development was comparable between genotypes and independent of diazoxide (Fig. 4 H). Therefore, both pharmacological inhibition of the NLRP3 inflammasome and pharmacological reduction of beta-cell secretory demand, attenuated the development of amyloid-induced hyperglycemia, supporting a model in which islet amyloid promotes beta-cell failure through NLRP3-dependent inflammation and increased secretory stress.

**Figure 4.**
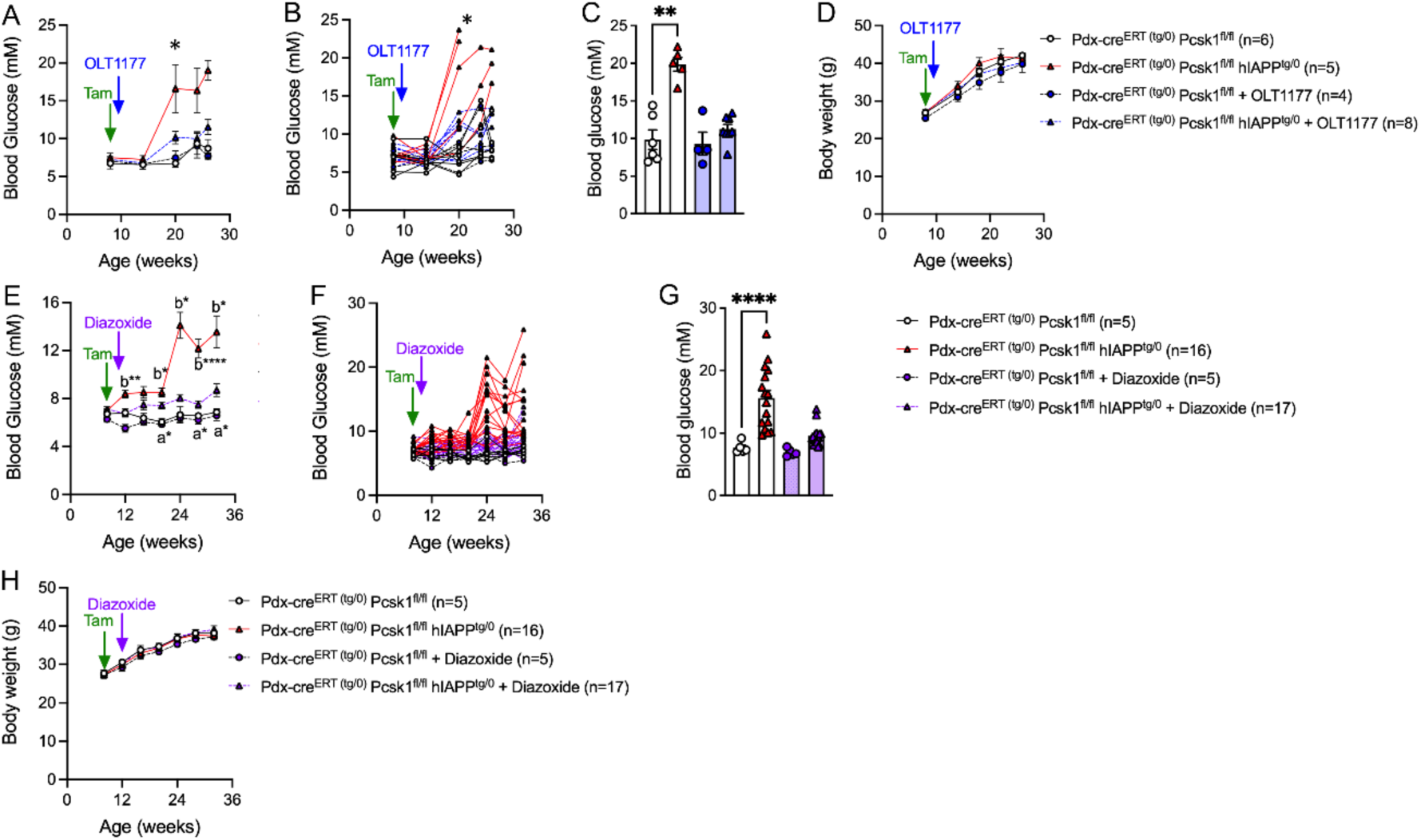
The NLRP3 inhibitor OLT1177 prevents the development of islet amyloid-induced hyperglycemia. (A) Average glycemia per experimental group and (B) glycemia of individual mice in Pdx-cre^ERT^ Pcsk1^fl/fl^ hIAPP^tg/0^ mice that received tamoxifen at 8 weeks of age and switched to a control diet or diet containing 7.5 g/kg OLT1177 at 9 weeks of age. * denotes a timepoint in which some animals were taken out of experiment due to reaching a humane endpoint (glycemia). (C) Highest blood glucose measurement reached during the study. (D) Body weight development in mice fed OLT1177 starting at 9 weeks of age. (E) Average glycemia per experimental group and (F) glycemia of individual mice in Pdx-cre^ERT^ Pcsk1^fl/fl^ hIAPP^tg/0^ mice that received tamoxifen at 8 weeks of age and switched to a control diet or diet containing 1g/kg diazoxide at 9 weeks of age. (G) Highest blood glucose measurement reached during the study. (H) Body weight development in mice fed diazoxide starting at 9 weeks of age. Statistics: A, B, D, E, F, H mixed-effects model. C, G: 1-way ANOVA with Tukey’s multi comparison test. * p<0.05, ** p< 0.01, ****p<0.0001.

## Discussion

In the present study, we present a mouse model that recapitulates key pathological features of human type 2 diabetes and we used this model to demonstrate that inhibition of the NLRP3 inflammasome prevents the development of severe hyperglycemia. By combining impaired proinsulin processing, the capacity to form islet amyloid deposits, and ageing, we generated a model that develops progressive beta-cell dysfunction, islet inflammation, loss of beta-cell mass, and overt diabetes. Importantly, these pathological changes were prevented by treatment with OLT1177, a selective oral NLRP3 inflammasome inhibitor.

Current preclinical models of type 2 diabetes rely predominantly on diet-induced obesity or genetic defects in leptin signaling. While these models have substantially advanced our understanding of metabolic disease, they incompletely reproduce the pathology observed in humans. Human type 2 diabetes is characterized by progressive beta-cell failure [7], impaired proinsulin processing [39–41] including reduced expression of islet prohormone convertase 1/3 in islets [45], and deposition of islet amyloid [9, 10, 19], pathological features that are largely absent from conventional rodent models. The resulting phenotype resembles advanced human type 2 diabetes more closely than currently available models and provides an experimental platform for investigating mechanisms of beta-cell failure and disease progression.

The absence of obesity in this mouse model is particularly relevant for the interpretation of our intervention studies. Several recently developed NLRP3 inhibitors have been reported to reduce food intake, body weight gain, and adiposity in obese rodents, effects that are accompanied by improvements in glycemia and other metabolic parameters [31, 32, 46]. While these findings support an important role of NLRP3 signaling in obesity-associated metabolic dysfunction, they make it difficult to distinguish direct anti-inflammatory effects from secondary consequences of weight loss. We therefore intentionally employed a non-obese model in which body weight was not different between groups and remained unaffected by OLT1177 treatment. The prevention of hyperglycemia observed in our study can therefore not be attributed to reduced adiposity or weight loss, but instead identifies a direct role of NLRP3-driven inflammation in the pathogenesis of beta-cell failure. This is also supported by another preclinical study, in which diet-induced obese mice that were also treated with the beta-cell toxin streptozotocin, showed reduced glycemia and increased circulating insulin levels after injection of OLT1177 [46].

Our findings further strengthen the concept that islet amyloid deposition is not merely a histological hallmark of type 2 diabetes but an active driver of disease progression. Human IAPP aggregates activate the NLRP3 inflammasome and stimulate IL-1beta production [6], thereby linking metabolic stress to innate immune activation. Consistent with this mechanism, amyloid deposition in our model was associated with increased macrophage accumulation, elevated expression of inflammatory cytokines including *Il1b* and *Il6*, and progressive loss of beta-cell mass. Previous studies demonstrated that blockade of IL-1 signaling improves glucose control and preserves beta-cell function in both experimental models and patients with type 2 diabetes [25–28]. As such, pharmacological blockage of IL-1 signaling in a mouse model of islet amyloidosis improved glycemia but did not change amyloid burden [26] while the same group reported reduced amyloid deposition in an islet transplant model [25] treated with a recombinant IL-1 receptor antagonist. Further, blocking IL-1beta signaling *in vitro* in isolated human islets reduced apoptosis and amyloid formation [47], suggesting that neutralizing IL-1beta dampens the interaction of immune cell-derived inflammation and beta-cell failure. Our results extend these observations upstream and provide direct evidence that activation of the NLRP3 inflammasome itself is a critical pathogenic event linking amyloid deposition to beta-cell dysfunction *in vivo*.

An important observation of the current study is the magnitude of the metabolic benefit achieved by NLRP3 inhibition. OLT1177 prevented the development of severe hyperglycemia and maintained glucose concentrations near those observed in mice unable to develop amyloid pathology. This degree of protection is remarkable given the aggressive nature of the model and suggests that inflammatory signaling is not simply a secondary consequence of beta-cell failure but contributes directly to disease progression. The partial protection observed with diazoxide supports the concept that inflammatory activation and increased secretory demand act in concert to promote beta-cell exhaustion. Together, these findings support a model in which amyloid-induced NLRP3 activation amplifies beta-cell stress through IL-1beta-dependent mechanisms, ultimately driving progressive beta-cell dysfunction and loss.

The translational relevance of these findings is supported by human data. Analysis of the subgroup of patients with type 2 diabetes in a double-blind placebo-controlled intervention study in patients with heart failure revealed a dose-dependent reduction in fasting plasma glucose with OLT1177 [38]. The greatest glucose reduction was observed at the highest dose tested and was evident after only two weeks of treatment. Although these analyses were exploratory and performed in a relatively small number of participants, the concordance between the clinical and experimental findings is notable. Together, the animal and human data provide converging evidence that NLRP3 inflammasome activation contributes to hyperglycemia in type 2 diabetes and that pharmacological inhibition of this pathway may offer therapeutic benefit.

The broader implications of these findings extend beyond OLT1177/dapansutrile itself. The NLRP3 inflammasome has emerged as an attractive therapeutic target, and multiple pharmaceutical companies have developed selective NLRP3 inhibitors that are currently being evaluated across a broad range of inflammatory and cardiometabolic disorders [29]. Our findings provide mechanistic evidence supporting the inclusion of type 2 diabetes among the indications most likely to benefit from this therapeutic strategy. Importantly, unlike many currently available glucose-lowering therapies that primarily compensate for metabolic dysfunction, NLRP3 inhibition targets a pathway implicated in the progressive loss of beta-cell function. This raises the possibility that suppression of inflammasome signaling may modify disease progression rather than merely improve glycemic control.

Several limitations should be acknowledged. The experimental intervention was initiated before the onset of advanced diabetes and therefore primarily addresses disease prevention rather than reversal of established beta-cell failure. In addition, although our data strongly support a role for NLRP3-dependent inflammation, the relative contribution of IL-1beta, IL-18, pyroptosis, and other downstream pathways of the NLRP3 inflammasome remains to be determined. Future studies will be required to establish whether NLRP3 inhibition can restore beta-cell function in established disease and to identify biomarkers predicting therapeutic response.

In conclusion, we describe a mouse model that captures key pathological hallmarks of human type 2 diabetes, including impaired proinsulin processing, islet amyloid deposition, inflammation, and progressive beta-cell failure. Using this model, we demonstrate that selective inhibition of the NLRP3 inflammasome prevents the development of overt diabetes independently of changes in body weight. Together with supportive human data [38], these findings provide compelling evidence that activation of the NLRP3 inflammasome is a central mechanism driving beta-cell failure in type 2 diabetes and support ongoing clinical efforts to evaluate selective NLRP3 inhibitors, including OLT1177/dapansutrile, as disease-modifying therapies for diabetes.

## Methods

### Mice

All animal experiments were performed in accordance with the federal laws of Switzerland including institutional guidelines and approved by the cantonal veterinary office of Basel. Pdx-cre^ERT^ Pcsk1^fl/fl^ mice on a C57BL/6N genetic background were described in detail before [43]. Briefly, Tg(Pdx1-Cre/Esr1*)^Dam^ (Jax: 024968) [48] were obtained from Michael B. Wheeler (University of Toronto) and crossed to B6N-Pcsk1^tm1Boe^ mice [49]. For the purpose of this project, mice were backcrossed to an in-house colony of FVB/NRj (Janvier, France) for at least 10 generations. Mice expressing human IAPP in their beta cells (Rip-hIAPP) [42] were obtained from Jackson Laboratories (Jax stain: 008232), backcrossed to our in-house FVB/NRj genetic background for at least 10 generations and then intercrossed with Pdx-cre^ERT^ Pcsk1^fl/fl^ on the FBV/NRj genetic background. Mice were housed in groups of 2–5 mice in individually-ventilated cages (Indulab, Gams, Switzerland) with standard bedding, nesting material, and a cardboard house, under specific pathogen-free conditions (FELASA), in a facility controlled for temperature (21 ± 2 °C) and humidity (50–60%) with a 12 h dark/12 h light cycle (lights on at 6:00), at the University of Basel. Mice received standard chow (3436, Kliba Nafag, Kaiseraugst, Switzerland). Where indicated, mice received Purified Open Standard Diet (D21051704i, Research Diets, New Brunswick, NJ, USA) containing 5 g/kg natural peanut powder with or without 7.5 g/kg OLT1177. OLT1177 was provided by Olatec (New York, USA). Where indicated, mice received standard chow containing 1 g/kg diazoxide (Proglicem, MSD, Munich, Germany). Diazoxide was mixed with powdered food, reaggregated into pellets and dried in an oven. Where indicated, mice were given tamoxifen (HY-13757A, MCE, Lucerna Chem, Lucerne, Switzerland) dissolved in ethanol/corn oil (C8267, Sigma-Aldrich, St Louis, MO, USA), 125 mg/kg body weight on Monday, Wednesday, and Friday at 8 weeks of age by oral gavage. Whenever possible, experimenters were blinded to the genotype of the experimental animal.

### Body weight development, food intake and glycemia

Body weight was measured by a Scout STX balance (Ohaus, Nänikon, Switzerland) using the dynamic weighing feature (average over 5 seconds). Food intake was assessed by measurement of the food given/left in the rack feeder of the cage. Blood glucose was measured from the tail vein using a glucometer (FreeStyle Freedom Lite, Abbott, Baar, Switzerland).

### Islet isolation, RNA isolation and real-time PCR

Pancreatic islets were isolated as described in detail before [50]. Briefly, collagenase (4189, Worthington, Lakewood, NJ, USA) was injected through the bile duct and pancreata were incubated at 37 °C for 34 minutes with occasional shaking. HBSS medium containing 0.5 w/v BSA was used to quench the reaction. Islets were hand-picked and resuspended in RPMI-1640 medium containing 11.1 mM glucose, 100 units/ml penicillin, 100 μg/ml streptomycin, 2 mM glutamax, 50 μg/ml gentamycin, 10 μg/ml fungizone and 10% FCS. RNA was isolated using the NucleoSpin RNA Kit (740955, Macherey Nagel, Oensingen, Switzerland). Reverse transcription was done with GoScript reverse transcriptase (A2801, Promega, Dübendorf, Switzerland) and real time qPCR with an ABI 7500 Fast Real-Time PCR System (Thermo Fisher Scientific, Waltham, MA, USA) with TaqMan assays (Thermo Fisher Scientific).

### Histological analyses

Pancreata were dissected, weighed, and incubated in 4% buffered formalin (Forma Fix, Hittnau, Switzerland) overnight at 4 °C, processed on a TPC 15 DUO, and embedded in paraffin on a TES Valida (Medite, Dietikon, Switzerland) and sectioned into 5 μm slices on an HM340E microtome (Thermo Fisher Scientific). Tissue sections were deparaffinized, rehydrated, and incubated with an antibody raised against insulin (181547, Abacam, UK) and CD45 to visualize lymph nodes (103102, Biolegend, USA). Slides were also incubated in 0.5% w/v Thioflavin S (Sigma-Aldrich) to visualize amyloid deposits and DAPI to visualize nuclei. For determination of beta-cell mass, tissue sections from three to four different depths of the pancreas (at least 100 μm apart) were stained for insulin, imaged using an automated Eclipse Ni microscope (Nikon, Egg, Switzerland) with an Orca Flash 4.0 camera (Hamamatsu Photonics, Solothurn, Switzerland), a 4× Objective NA 0.2 (Nikon), and a PL-200 slide loading robot (PRIOR Scientific, Fulbourn, UK) using NIS Elements software v5. Pancreas and islet morphology was quantified with QuPath [51]. Beta-cell mass was calculated as pancreas weight × sum (insulin-positive area)/sum (pancreas area). For studies quantifying amyloid and insulin areas, Image J Fiji [52] was used. Amyloid severity is thioflavin S-positive area / islet area x 100.

Amyloid prevalence is the number of thioflavin S-positive islets / total number of islets x 100. Insulin severity is insulin-positive area / islet area x 100. For quantification of macrophages, sections were incubated overnight at 4°C with rat anti-Mac2 (CL8942AP, Cederlane) primary antibody followed the next day with anti-rat HRP (A10549, Invitrogen) secondary antibody for 2 hours at room temperature. Washing steps were performed using PBS containing Tween 0.05%. Signal detection was performed using 3,3-diaminobenzidine (DAB; DAKO Cat# K3468) HRP substrate, and sections were mounted under coverslips using Pertex mounting medium (00840, Histolab). Slides were scanned using an Olympus VS200 with a 20X objective. Islet pictures were selected and analyzed using QuPath. Intra-islet and peri-islet Mac2^+^ macrophages were manually quantified, with peri-islet macrophages defined as cells touching the islet boundary.

### Flow cytometry

Blood was collected from the tail vain of mice and stored in EDTA-coated tubes at room temperature until further processing. Subsequently, samples were incubated for 10 min in CD16/32 FcBlock (1:100, BD Biosciences #553142) in flow cytometry buffer (PBS supplemented with 2% FBS and 50 mM EDTA) at room temperature. After, the following murine antibodies were added: CD45-Alexa Fluor700 (1:200, BioLegend #103128), CD11b-PE (1:200, BioLegend #101208), Ly6G-APC-Cy7 (1:400, BioLegend #127624), CD3-PerCP-Cy5.5 (1:100, BioLegend #100218) for 30 min at room temperature in the dark, diluted at corresponding dilutions in flow cytometry buffer. Samples were then lysed and fixed using the BD FACS Lysing solution (BD Biosciences #349202) until samples became transparent. Subsequently, samples were washed twice with flow cytometry buffer by centrifuging at 600 g for 5 min and discarding the supernatant. Sample acquisition was performed on a 3-Laser Cytoflex (Beckman Coulter) with a subsequent data analysis in FlowJo v10.10.0 (BD Biosciences). Gating strategy is depicted in Fig. S2I.

### Software

The manuscript was prepared using Microsoft Office version 16.

## Statistical analyses

Data are presented as mean ± SEM. The statistical test used are indicated in the figure legends. Statistical analyses were conducted with Prism v11 (GraphPad Software, San Diego, CA, USA).

## Acknowledgements

We thank the animal facilities and services of the University of Basel for animal husbandry and the Department of Biomedicine histology core facility as well as the microscopy core facility for support and for providing infrastructure. We acknowledge Joyce de Paula Souza and Hélène Mereau (University of Basel) for technical assistance. A.L.G. was supported by Horizon Europe (Intercept-t2D, 101095433). M.Y.D is supported by the Swiss National Science Foundation (214900 and 226049) and the European Union – Horizon/Swiss State Secretariat for Education, Research and Innovation (101095433).

## Authors’ relationships and activities

Olatec (New York, USA) provided food containing OLT1177 for the current study but did not finance or influence the study in any way. M.Y.D. is part of the clinical advisory board of Olatec and receives OLT1177 for an ongoing clinical study funded by a European Union/Swiss grant.

---

**Figure S1.**
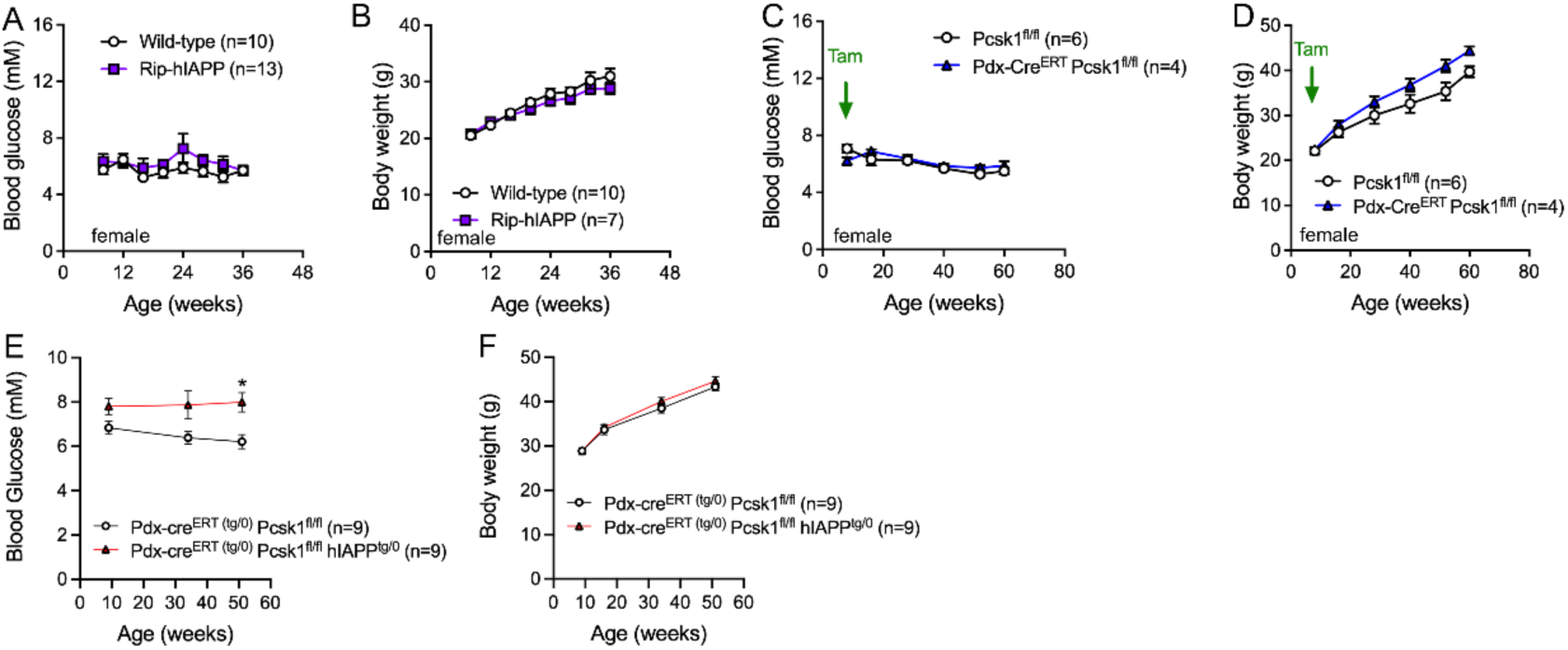
Body weight and glycemia of individual mouse lines. (A) Glycemia and (B) body weight development in female hIAPP mice on an FVB/N genetic background. (C) Glycemia and (D) body weight development in female Pdx-cre^ERT^ Pcsk1^fl/fl^ mice that had received tamoxifen at 8 weeks of age. (E) Glycemia and (F) body weight development in male Pdx-cre^ERT^ Pcsk1^fl/fl^ hIAPP^tg/0^ mice without tamoxifen application. Statistics: A-F 2-way ANOVA with Sidak’s multi comparison test. * p<0.05.

**Figure S2.**
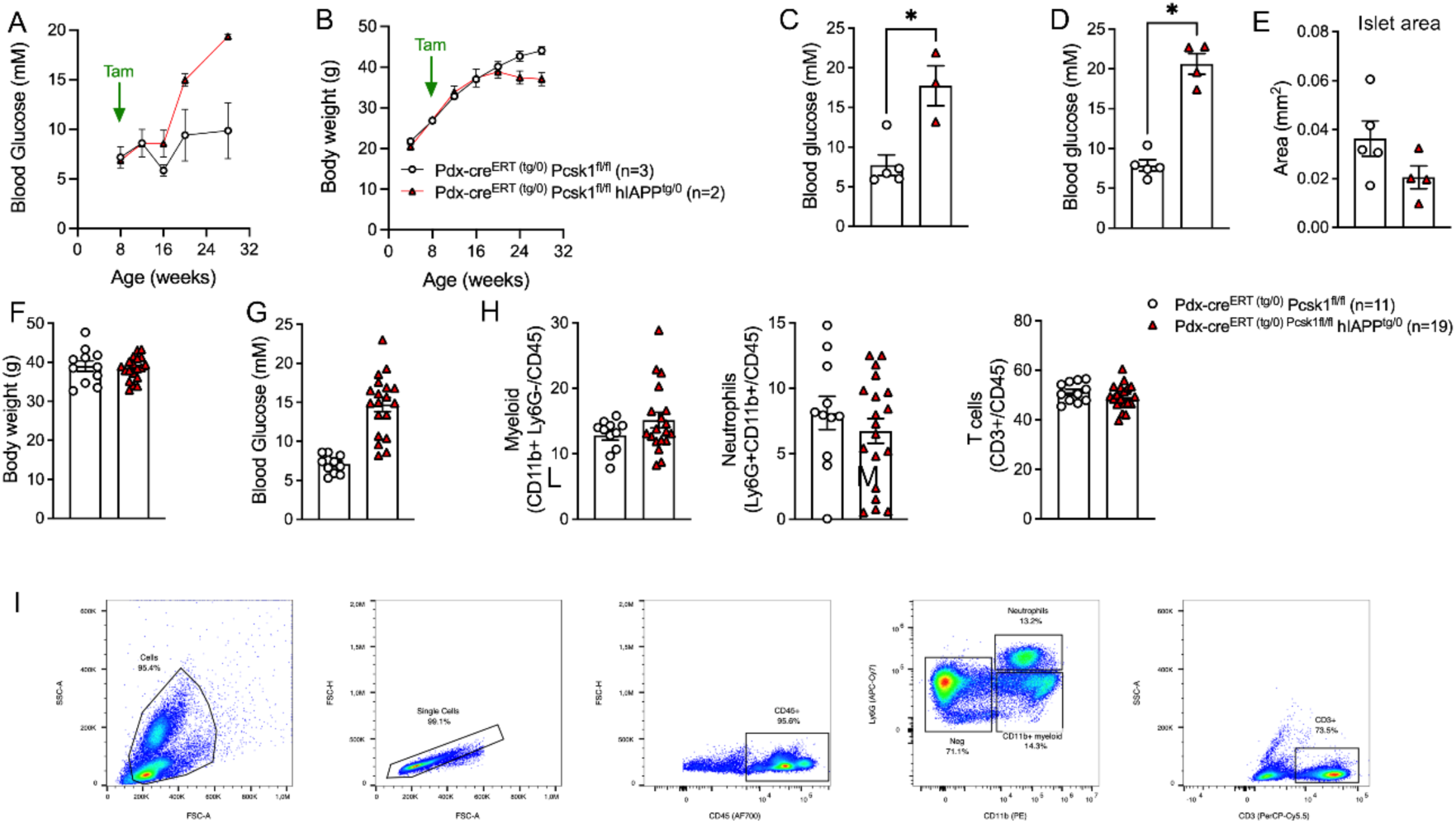
Additional data of Pdx-cre^ERT^ Pcsk1^fl/fl^ hIAPP^tg/0^ mice. (A) Glycemia and (B) body weight development in a cohort of Pdx-cre^ERT^ Pcsk1^fl/fl^ hIAPP^tg/0^ mice used for histological analysis. (C) Blood glucose values at sacrifice in a cohort of 28-week-old Pdx-cre^ERT^ Pcsk1^fl/fl^ hIAPP mice used for islet isolation. (D) Glycemia and (E) islet area per section in a cohort of 28-week-old Pdx-cre^ERT^ Pcsk1^fl/fl^ hIAPP mice used for macrophage quantification. (F) Body weight at sacrifice, (G) blood glucose values at sacrifice and (H) flow cytometry analysis of blood collected from 20-28-week-old Pdx-cre^ERT^ Pcsk1^fl/fl^ hIAPP^tg/0^ mice. (I) Corresponding gating strategy. Statistics: C-H: Mann-Whitney test. * p<0.05.

## References

1. Martinon, F., K. Burns, and J. Tschopp, The inflammasome: a molecular platform triggering activation of inflammatory caspases and processing of proIL-beta. Mol Cell, 2002. 10(2): p. 417–26.

2. Zhou, R., et al., Thioredoxin-interacting protein links oxidative stress to inflammasome activation. Nat Immunol, 2010. 11(2): p. 136–40.

3. Martinon, F., et al., Gout-associated uric acid crystals activate the NALP3 inflammasome. Nature, 2006. 440(7081): p. 237–41.

4. Tschop, M. and G. Thomas, Fat fuels insulin resistance through Toll-like receptors. Nat Med, 2006. 12(12): p. 1359–61.

5. Duewell, P., et al., NLRP3 inflammasomes are required for atherogenesis and activated by cholesterol crystals. Nature, 2010. 464(7293): p. 1357–61.

6. Masters, S.L., et al., Activation of the NLRP3 inflammasome by islet amyloid polypeptide provides a mechanism for enhanced IL-1beta in type 2 diabetes. Nat Immunol, 2010. 11(10): p. 897–904.

7. Rohm, T.V., et al., Inflammation in obesity, diabetes, and related disorders. Immunity, 2022. 55(1): p. 31–55.

8. Kahn, S.E., et al., Evidence of cosecretion of islet amyloid polypeptide and insulin by beta-cells. Diabetes, 1990. 39(5): p. 634–8.

9. Westermark, P., Quantitative studies on amyloid in the islets of Langerhans. Ups J Med Sci, 1972. 77(2): p. 91–4.

10. Jurgens, C.A., et al., beta-cell loss and beta-cell apoptosis in human type 2 diabetes are related to islet amyloid deposition. Am J Pathol, 2011. 178(6): p. 2632–40.

11. Hull, R.L., et al., Islet amyloid: a critical entity in the pathogenesis of type 2 diabetes. J Clin Endocrinol Metab, 2004. 89(8): p. 3629–43.

12. Westermark, P., et al., Islet amyloid polypeptide: pinpointing amino acid residues linked to amyloid fibril formation. Proc Natl Acad Sci U S A, 1990. 87(13): p. 5036–40.

13. Lorenzo, A., et al., Pancreatic islet cell toxicity of amylin associated with type-2 diabetes mellitus. Nature, 1994. 368(6473): p. 756–60.

14. Maedler, K., et al., Glucose-induced beta cell production of IL-1beta contributes to glucotoxicity in human pancreatic islets. J Clin Invest, 2002. 110(6): p. 851–60.

15. Hou, X., et al., Prolonged exposure of pancreatic beta cells to raised glucose concentrations results in increased cellular content of islet amyloid polypeptide precursors. Diabetologia, 1999. 42(2): p. 188–94.

16. Eguchi, K., et al., Saturated fatty acid and TLR signaling link beta cell dysfunction and islet inflammation. Cell Metab, 2012. 15(4): p. 518–33.

17. Ehses, J.A., et al., Increased number of islet-associated macrophages in type 2 diabetes. Diabetes, 2007. 56(9): p. 2356–70.

18. Boni-Schnetzler, M. and D.T. Meier, Islet inflammation in type 2 diabetes. Semin Immunopathol, 2019. 41(4): p. 501–513.

19. Clark, A., et al., Islet amyloid, increased A-cells, reduced B-cells and exocrine fibrosis: quantitative changes in the pancreas in type 2 diabetes. Diabetes Res, 1988. 9(4): p. 151–9.

20. Meier, D.T., et al., Islet amyloid formation is an important determinant for inducing islet inflammation in high-fat-fed human IAPP transgenic mice. Diabetologia, 2014. 57(9): p. 1884–8.

21. Park, Y.J., et al., Dual role of interleukin-1beta in islet amyloid formation and its beta-cell toxicity: Implications for type 2 diabetes and islet transplantation. Diabetes Obes Metab, 2017. 19(5): p. 682–694.

22. Stienstra, R., et al., The inflammasome-mediated caspase-1 activation controls adipocyte differentiation and insulin sensitivity. Cell Metab, 2010. 12(6): p. 593–605.

23. Vandanmagsar, B., et al., The NLRP3 inflammasome instigates obesity-induced inflammation and insulin resistance. Nat Med, 2011. 17(2): p. 179–88.

24. Wen, H., et al., Fatty acid-induced NLRP3-ASC inflammasome activation interferes with insulin signaling. Nat Immunol, 2011. 12(5): p. 408–15.

25. Westwell-Roper, C., et al., IL-1 blockade attenuates islet amyloid polypeptide-induced proinflammatory cytokine release and pancreatic islet graft dysfunction. J Immunol, 2011. 187(5): p. 2755–65.

26. Westwell-Roper, C.Y., et al., IL-1 mediates amyloid-associated islet dysfunction and inflammation in human islet amyloid polypeptide transgenic mice. Diabetologia, 2015. 58(3): p. 575–85.

27. Larsen, C.M., et al., Interleukin-1-receptor antagonist in type 2 diabetes mellitus. N Engl J Med, 2007. 356(15): p. 1517–26.

28. Kataria, Y., C. Ellervik, and T. Mandrup-Poulsen, Treatment of type 2 diabetes by targeting interleukin-1: a meta-analysis of 2921 patients. Semin Immunopathol, 2019. 41(4): p. 413–425.

29. Vande Walle, L. and M. Lamkanfi, Drugging the NLRP3 inflammasome: from signalling mechanisms to therapeutic targets. Nat Rev Drug Discov, 2024. 23(1): p. 43–66.

30. Meier, D.T., J. de Paula Souza, and M.Y. Donath, Targeting the NLRP3 inflammasome-IL-1beta pathway in type 2 diabetes and obesity. Diabetologia, 2025. 68(1): p. 3–16.

31. Bultinck, J., et al., NLRP3 inhibition by VTX3232 tempers inflammation resulting in reduced body weight, hyperglycemia, and hepatic steatosis in obese male mice. Mol Metab, 2026. 103: p. 102282.

32. Thornton, P., et al., Reversal of High Fat Diet-Induced Obesity, Systemic Inflammation, and Astrogliosis by the NLRP3 Inflammasome Inhibitors NT-0249 and NT-0796. J Pharmacol Exp Ther, 2024. 388(3): p. 813–826.

33. Marchetti, C., et al., OLT1177, a beta-sulfonyl nitrile compound, safe in humans, inhibits the NLRP3 inflammasome and reverses the metabolic cost of inflammation. Proc Natl Acad Sci U S A, 2018. 115(7): p. E1530–E1539.

34. Marchetti, C., et al., NLRP3 inflammasome inhibitor OLT1177 suppresses joint inflammation in murine models of acute arthritis. Arthritis Res Ther, 2018. 20(1): p. 169.

35. Toldo, S., et al., The NLRP3 Inflammasome Inhibitor, OLT1177 (Dapansutrile), Reduces Infarct Size and Preserves Contractile Function After Ischemia Reperfusion Injury in the Mouse. J Cardiovasc Pharmacol, 2019. 73(4): p. 215–222.

36. Sanchez-Fernandez, A., et al., OLT1177 (Dapansutrile), a Selective NLRP3 Inflammasome Inhibitor, Ameliorates Experimental Autoimmune Encephalomyelitis Pathogenesis. Front Immunol, 2019. 10: p. 2578.

37. Kluck, V., et al., Dapansutrile, an oral selective NLRP3 inflammasome inhibitor, for treatment of gout flares: an open-label, dose-adaptive, proof-of-concept, phase 2a trial. Lancet Rheumatol, 2020. 2(5): p. e270–e280.

38. Wohlford, G.F., et al., Phase 1B, Randomized, Double-Blinded, Dose Escalation, Single-Center, Repeat Dose Safety and Pharmacodynamics Study of the Oral NLRP3 Inhibitor Dapansutrile in Subjects With NYHA II-III Systolic Heart Failure. J Cardiovasc Pharmacol, 2020. 77(1): p. 49–60.

39. Kahn, S.E., et al., Proinsulin as a marker for the development of NIDDM in Japanese-American men. Diabetes, 1995. 44(2): p. 173–9.

40. Kahn, S.E. and P.A. Halban, Release of incompletely processed proinsulin is the cause of the disproportionate proinsulinemia of NIDDM. Diabetes, 1997. 46(11): p. 1725–32.

41. Mykkanen, L., et al., Serum proinsulin levels are disproportionately increased in elderly prediabetic subjects. Diabetologia, 1995. 38(10): p. 1176–82.

42. Janson, J., et al., Spontaneous diabetes mellitus in transgenic mice expressing human islet amyloid polypeptide. Proc Natl Acad Sci U S A, 1996. 93(14): p. 7283–8.

43. Meier, D.T., et al., Prohormone convertase 1/3 deficiency causes obesity due to impaired proinsulin processing. Nat Commun, 2022. 13(1): p. 4761.

44. Dror, E., et al., Postprandial macrophage-derived IL-1beta stimulates insulin, and both synergistically promote glucose disposal and inflammation. Nat Immunol, 2017. 18(3): p. 283–292.

45. Fadista, J., et al., Global genomic and transcriptomic analysis of human pancreatic islets reveals novel genes influencing glucose metabolism. Proc Natl Acad Sci U S A, 2014. 111(38): p. 13924–9.

46. Wu, Y., et al., Dapansutrile preserves pancreatic beta-cell function by regulating insulin vesicle membrane fusion and mitochondrial homeostasis. Diabetes Obes Metab, 2026. 28(4): p. 3054–3069.

47. Hui, Q., et al., Amyloid formation disrupts the balance between interleukin-1beta and interleukin-1 receptor antagonist in human islets. Mol Metab, 2017. 6(8): p. 833–844.

48. Gu, G., J. Dubauskaite, and D.A. Melton, Direct evidence for the pancreatic lineage: NGN3+ cells are islet progenitors and are distinct from duct progenitors. Development, 2002. 129(10): p. 2447–57.

49. Traub, S., et al., Pancreatic alpha Cell-Derived Glucagon-Related Peptides Are Required for beta Cell Adaptation and Glucose Homeostasis. Cell Rep, 2017. 18(13): p. 3192–3203.

50. Steiger, L., et al., Protocol for isolation and spectral flow cytometry analysis of immune cells from the murine exocrine and endocrine pancreas. STAR Protoc, 2023. 4(4): p. 102664.

51. Bankhead, P., et al., QuPath: Open source software for digital pathology image analysis. Sci Rep, 2017. 7(1): p. 16878.

52. Schindelin, J., et al., Fiji: an open-source platform for biological-image analysis. Nat Methods, 2012. 9(7): p. 676–82.

